# Are brain displacements and pressures within the parenchyma induced by surface pressure differences? A computational modelling study

**DOI:** 10.1101/2022.09.07.506967

**Authors:** Eleonora Piersanti, Marie E. Rognes, Vegard Vinje

## Abstract

The intracranial pressure is implicated in many homeostatic processes in the brain and is a fundamental parameter in several diseases such as e.g. idiopathic normal pressure hydrocephalus (iNPH). The presence of a small but persistent pulsatile intracranial pulsatile transmantle pressure gradient (on the order of a few mmHg/m at peak) has recently been demonstrated in iNPH subjects. A key question is whether pulsatile ICP and displacements can be induced by a small pressure gradient originating from the brain surface e.g. pial arteries alone. In this study, we model the brain parenchyma as either a linearly elastic or a poroelastic medium and impose a pulsatile pressure gradient acting between the ventricular and the pial surfaces. Using this high-resolution physics-based model, we compute the effect of the pulsatile pressure gradient on parenchyma displacement, volume change, fluid pressure, and fluid flux. The resulting displacement field is pulsatile and in qualitatively and quantitatively good agreement with the literature, both with elastic and poroelastic models. However, the pulsatile forces on the boundaries are not sufficient for pressure pulse propagation through the brain parenchyma. Our results suggest that pressure differences originating over the brain surface via e.g. pial artery pulsatility are not sufficient to drive interstitial fluid (ISF) flow within the brain parenchyma and that potential pressure gradients found within the parenchyma rather arise from local pressure pulsations of blood vessels within the brain parenchyma itself.

## Introduction

The cerebrospinal fluid (CSF) is mainly contained in the subarachnoid space (SAS) and subarachnoid cisterns surrounding the brain parenchyma [1] and plays an important role in maintaining the homeostasis of the brain [2]. Intracranial pressure (ICP), both its static and pulsatile components, is involved in many of these homeostatic processes [3]. Its fluctuations are related to blood flow, respiration, and cerebral spinal fluid (CSF) flow in the brain. ICP has been the subject of investigations for many years and it is a fundamental parameter to diagnose diseases such as idiopathic normal pressure hydrocephalus (iNPH), and other forms of hydrocephalus [4]. The ICP can also be influenced by changes in anatomy, obstruction of the aqueduct, and traumatic brain injuries for example, and by changes in the material properties of the brain parenchyma due to ageing. In hydrocephalus and iNPH, the intracranial pressure pulsations increase [5], and moreover shunt response may be predicted by the pre-surgical pulse pressure [6].

The presence of a transmantle pressure gradient between different areas of the brain has been controversial. Some studies have reported the absence of a transmantle gradient in iNPH patients [7,8], and therefore excluded it among the causes of iNPH. Moreover, a study from Eide [9] showed how an uneven distribution of intracranial pulsatility is found in hydrocephalus patients. In more recent work from Vinje et al. [10] a small pulsatile gradient was analysed and quantified between the subdural and intraventricular ICP. The analysis of [10] is based on overnight intracranial pressure measurements from subarachnoid and ventricles areas, in 10 iNPH patients. The pulsatile gradient is mainly characterized by a cardiac component of mean amplitude 1.46 mmHg/m and a respiratory component of mean amplitude 0.52 mmHg/m. However, to what extent these gradients affect pulsatile brain displacements has not yet been investigated. Moreover, the mechanisms behind this pulsatile gradient are not fully understood.

Pressure pulsations within the parenchyma may have at least two possible origins: a “systemic” pressure pulse propagating via the surrounding subarachnoid space (SAS) and then travelling through the entire brain tissue, or from blood vessel pulsations distributed with the parenchyma. The question is ultimately whether the pressure pulse travels through the brain tissue (extracellular matrix) or the blood vessels (blood vessel network). The answer is relevant to assess the possibility of perivascular flow also within the brain (along arterioles and capillaries), as pulsations originating from blood vessels have been suggested to drive bulk flow of perivascular fluid [11–13]. In addition, cardiovascular pulsations in the brain can be linked to Alzheimer’s disease: cardiovascular impulse latency, propagation speed and direction present very different behaviour in patients with Alzheimer’s disease and age-matched healthy volunteers [14].

The scope of the present paper is therefore to investigate the origin and effects of intracranial pressure gradient pulsatility on the brain parenchyma. We consider two different models for the brain parenchyma: linear elasticity and a one–network poroelasticity (Biot). The pulsatile pressure gradient is modelled as in [10] and it is applied to each model via appropriate boundary conditions. For the poroelastic case(s), we model the pial surface as permeable or as impermeable. For the linear elasticity case, we study the effect of the cardiac component for the pulsatile gradient alone, and the effect of brain parenchyma incompressibility. In this study, we observed a pulsating displacement field both with the linear elasticity and the poroelasticity model. With the poroelasticity model, we observe that the pressure prescribed on the pial surface does not propagate into the parenchyma neither with the impermeable nor a permeable pial membrane. With the current parameters and boundary conditions, we conclude that it is unlikely that a pulsating pressure difference between the pial and ventricular boundaries is responsible for fluid pressure propagation or interstitial fluid flow in the parenchyma.

## Materials and methods

### Computational domain

The computational domain is based on the Colin27 human adult brain atlas FEM mesh [15](Figure 1a). This mesh consists of gray and white matter regions. Boundary markers were created to divide the domain boundary into one pial and one ventricular surface. The ventricular boundary is shown in Figure 1b. The mesh consists of 1 227 992 cells and 265 085 vertices. The minimum cell size *h_min_* is 0.1 mm and the maximum cell size *h_max_* is 15.7 mm

**Fig 1.**
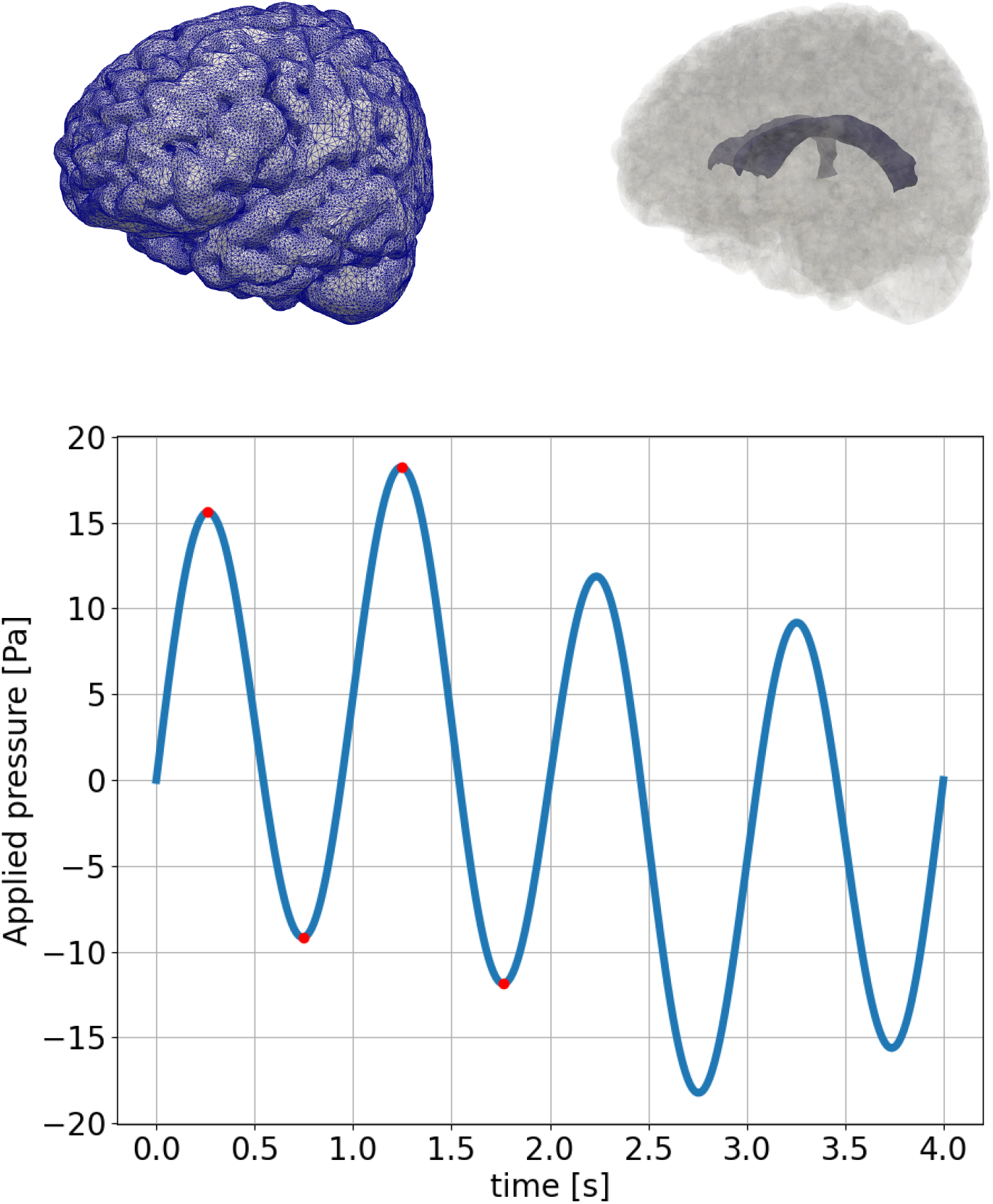
The computational mesh with edges (top left). The computational domain with highlighted ventricular surface (top right). The time-dependent applied pressure difference between the pial and ventricular surface (bottom). The red dots (*t* = 0.2625, 0.75, 1.25, 1.75) represent point of interest (peaks and valleys for *t* < 2.0).

### Governing equations

#### Linear elasticity

We first model the deformation of the brain parenchyma as that of an isotropic elastic solid as follows: find the displacement *u* such that

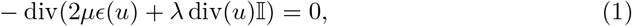

where *u* = *u*(*x, t*), for *x* ∈ Ω ⊂ ℝ^3^, *t* ∈ (0, *T*], and 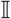 is the 3 × 3 identity matrix. The parameters λ > 0 and *μ* > 0 are the Lamé elasticity constants.

#### Poroelasticity

Biot’s equations describe a linear, isotropic solid permeated by a single fluid network. The equations read as follows: find the displacement *u* = *u*(*x,t*) and the fluid pressure *p* = *p*(*x, t*) such that

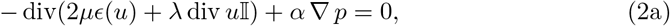

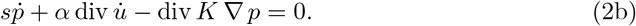

In addition to the parameters present in the linear elasticity system (1), we define the Biot-Willis coefficient *α* ∈ (0,1], the storage coefficient *s* > 0, and the hydraulic conductivity tensor *K* = *κ/v* > 0 with *κ* and *v* being the permeability and fluid viscosity, respectively. The (Darcy) fluid velocity *v*, representing the fluid velocity within the porous network i.e. the flow of interstitial fluid, is defined as

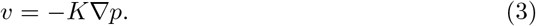

### Boundary conditions

We consider numerical experiments with different sets of boundary conditions for the elasticity and poroelasticity models. Specifically, we set a pressure difference between the subarachnoid space (pial surface) and the ventricles (ventricular surface). The pressure gradient is modelled, using data from [10], as a combination of two sinusoidal functions representing the cardiac cycle, with period *T_c_* = 1 s, and the respiratory cycle, with period of *T_r_* = 4s, respectively. With coefficients *a_c_* = 1.46 mmHg/m, *a_r_* = 0.52 mmHg/m, and assuming a brain width of *L* = 7 cm, we then compute the pressure difference between the pial and ventricular surface as d*p* = (*a_c_* sin(2*πt*) + *a_r_* sin(0.5*πt*))*L*. Specifically, we set the pressure gradient (Fig. 1) as

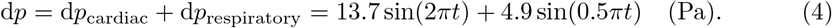

#### Linear elasticity

We impose a no-stress condition on the ventricular surface and a time-dependent pressure on the pial boundary of the parenchyma, resulting in the prescribed pressure difference dp. First, we investigate the effect of the cardiac cycle alone (Model A cf. Table 2), thus imposing the following Neumann-type boundary conditions

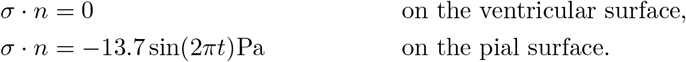

**Table 1.** Overview of material parameters used in numerical simulations, values and literature references.

| Name | Symbol | Values | Units | Reference |
| --- | --- | --- | --- | --- |
| Poisson ratio | $\nu$ | 0.495, 0.4983 | [-] | [17, 18] |
| Young's modulus | $E$ | 1642 | Pa | [19] |
| Biot-Willis coefficient | $\alpha$ | 1.0 | [-] | [20] |
| Hydraulic conductivity | $K$ | $1.57 \cdot 10^{-5}$ | $\text{mm}^2 \text{Pa}^{-1} \text{s}^{-1}$ | [16] |
| Storage coefficient | $s$ | $3.9 \cdot 10^{-4}$ | $\text{Pa}^{-1}$ | [20] |
| Pial permeability | $\beta$ | $0, \infty$ | $\text{mmPa}^{-1} \text{s}^{-1}$ | |

**Table 2.** Overview of computational models

| Model | Constitutive equations | Forces | $\nu$ | $\beta$ |
| --- | --- | --- | --- | --- |
| A | Elastic | Cardiac | 0.495 | NA |
| B | Elastic | Cardiac and respiratory | 0.495 | NA |
| C | Elastic | Cardiac and respiratory | 0.4983 | NA |
| D | Poroelastic | Cardiac and respiratory | 0.495 | 0 |
| E | Poroelastic | Cardiac and respiratory | 0.495 | ∞ |

**Table 3.** The mesh used for the mesh convergence: name, number of cells, minimum cell diameter and maximum cell diameter.

| Name | number of cells | $h_{min}$ | $h_{max}$ |
| --- | --- | --- | --- |
| COARSE | 99605 | 0.49 | 26.74 |
| COARSE-GR1 | 796840 | 0.25 | 16.7386 |
| COARSE-GR2 | 6374720 | 0.10 | 8.70 |
| COARSE-RV | 153499 | 0.25 | 25.1336 |
| COARSE-RV-GR | 1227992 | 0.10 | 15.6639 |
| COLIN27 | 249361 | 0.49 | 13.50 |
| COLIN27-GR1 | 1994888 | 0.20 | 9.18 |

Second, we consider the combined effect of the cardiac and respiratory cycle on the pressure difference (Model B-C cf. Table 2) with the following Neumann boundary conditions

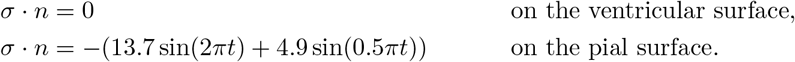

#### Poroelasticity

In the poroelastic model, we impose pure Neumann conditions for the total stress 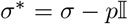:

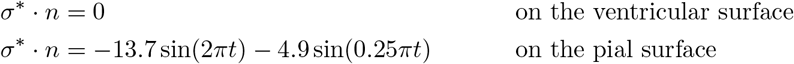

We introduce the parameter *β* ∈ [0, ∞) as the (membrane) permeability of the pial membrane and the additional boundary conditions

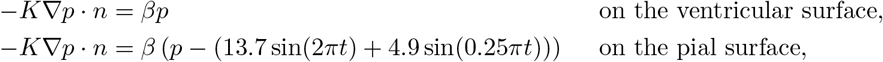

These Robin-type boundary conditions model the pia as a partially permeable membrane with *β* = (0, ∞), where for the lower bound *β* = 0 the pial membrane is impermeable (Neumann condition, Model D cf. Table 2), and for *β* → ∞ the pial membrane is fully permeable (Dirichlet condition, Model E cf. Table 2).

### Material parameters and model variations

We systematically consider a set of parameters (Tab 1) and of models (Tab 2). In models A, B, and C, the brain parenchyma is modelled as an elastic medium, while for models D and E the parenchyma is modeled as a poroelastic medium permeated by a single fluid network. We also investigate the effect of different external forces via boundary conditions, and of different parameters. For model A, the pulsatile pressure difference includes the cardiac component only while for the other models the pulsatile pressure difference is the combination of the cardiac and respiratory components. Regarding the elastic properties of the brain parenchyma, we consider a rather incompressible material *v* = 0.495 for models A, B, D, and E, and we investigate the effect of an even more incompressible material *v* = 0.4983 in model C. To model flow in the tortuous ECS in the poroelastic cases (D-E), we consider a small hydraulic conductivity [16] and we consider two different scenarios with a fully permeable or impermeable pial membrane.

### Numerical methods

The equations were solved with the finite element with FEniCS [21]. We used P1 elements for the displacement in linear elasticity (Models A,B,C) and the lowest-order Taylor-Hood elements (P2-P1) [22] for displacement and pressure (Models D, E). For all models, we impose a Neumann boundary condition for the momentum equation(s) on the entire boundary. This setup implies that the solutions are determined up to rigid motions or a constant pressure only, therefore we imposed additional constraints via Lagrange multipliers [23]. We used the implicit Euler scheme to discretize the equations in time. We performed convergence tests for several computational domains (see Supporting Information, Section B).

### Quantities of interest

For models A, B, and C, we report the displacement field *u*, its magnitude, and its values in selected points. The corresponding volume change is computed as:

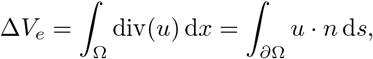

where we applied the divergence theorem. From the displacement field *u*, we compute the elastic stress tensor *σ*, and the von Mises stress *σ_M_* as:

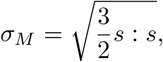

where *s* is the deviatoric part of the stress tensor defined as 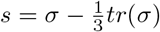. The von Mises stress provides information on the deviatoric stress while being a scalar value and therefore easier to visualize.

For the poroelastic models D and E, in addition to the above mentioned quantities, we analyse the fluid pressure *p* on the whole domain, and in selected planes and lines. The Darcy fluid velocity is computed as

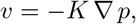

the fluid flux on the domain boundary is computed as

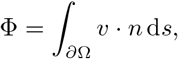

and the volume change caused by the fluid flux is

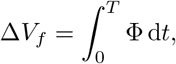

where *T* defines the time period of interest (e.g. *T* = 1s for the cardiac cycle). To compare flow and volume changes to values in the literature, we distinguish between volume changes occurring due to expansion and flow at the pial membrane and expansion and flow at the ventricular surface.

To quantify the stroke volumes caused by elastic deformation and by the fluid flow we consider the curves Δ*V_e_*, and Δ*V_f_* respectively. In particular, we identify the peak and valley in the time period of interest [0, *T*], sum their absolute values and divide by two. In the following, we describe the process to decompose the volume change curves into their cardiac and respiratory components.

### Separation of cardiac and respiratory components

The prescribed pulsatile pressure gradient is composed of a cardiac component (*T* = 1s) and a respiratory component (*T* = 4s). It is therefore natural to decompose the quantities of interest (described in the dedicated section) into the same components. First, consider the volume change caused by the elastic displacement. To compute the amplitude of the cardiac component of Δ*V_e_* (t) we compute the arithmetic average of the magnitude of peaks and valleys of Δ*V_e_* function over the 4s simulation for a total of 8 data points. Therefore, the cardiac component can be expressed as

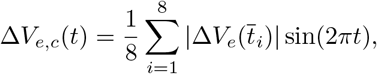

where 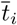 are the times corresponding to peaks or valleys of the function Δ*V_e_* (t). The corresponding respiratory component can be obtained by subtraction as follows

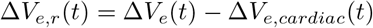

The above operations can be repeated on the volume change curves of pial and ventricles separately.

Similarly, the fluid flux can be decomposed into its cardiac and respiratory components. However, these fluctuations are not expected to be in phase with the pressure pulsations (as is the case for displacements). In this case, we therefore first identify the amplitude of the respiratory component. To this end, we impose that the value of the respiratory component Φ_*r*_ at t = 1s is the average between the peak value and the valley value in the neighbourhood of the chosen time t = 1s. Therefore the respiratory component can be expressed as

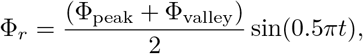

where Φ_peak_ and Φ_valley_ are the peak and valley values in the neighbourhood of *t* = 1s. The cardiac component can be obtained by subtraction

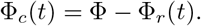

The above operations can be repeated on the flux curves of pial and ventricles separately.

### Mesh convergence test

We performed numerical convergence tests using several meshes. All the mesh refinings were performed in the FEniCS software. From the Colin 27 mesh [15] (COLIN27), we generated a finer mesh (COLIN27-GR1). From a coarsened version of the COLIN27 (COARSE) we generated two refined meshes: COARSE-GR1 and COARSE-GR2 where we applied the global refinement function in FEniCS once and twice, respectively. From the coarsened mesh COARSE, we also derived a mesh locally refined around the ventricular area (COARSE-RV), targeting the cells whose distance from the ventricles center *d* was *d* < 30 mm. Again, we performed a global refinement of the mesh COARSE-RV to obtain COARSE-RV-GR1. For the meshes described above, and listed with further details in 3, we simulated the linear elasticity equations described in 1 with *P*_1_ finite element for the displacement u and the following pure Neumann boundary condition for the total stress such as

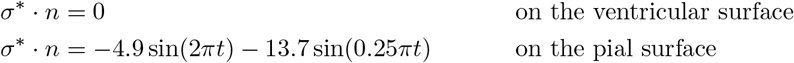

We then compared the volume change of the pial and ventricular surfaces for the different meshes as shown in Fig 2. Meshes COARSE-RV-GR and COLIN27-GR1 yield very similar results: the max values differences are 1% for the pial and 1.5% for the ventricular volume change. The COARSE-RV-GR is less computationally expensive since it has 1 227 992 cells compared to 1 994 888 cells of COLIN27-GR1. The maximal difference in the quantities computed on COARSE-RV-GR and COARSE-GR2 is 12% for the change of volume at the ventricular surface, and 6.47% for the change of volume at the pial surface. In addition, COARSE-GR2 contains 5.2 times the number of cells of COARSE-RV-GR, making it computationally expensive. Therefore, we chose to perform our computations on mesh COARSE-RV-GR.

**Fig 2.**
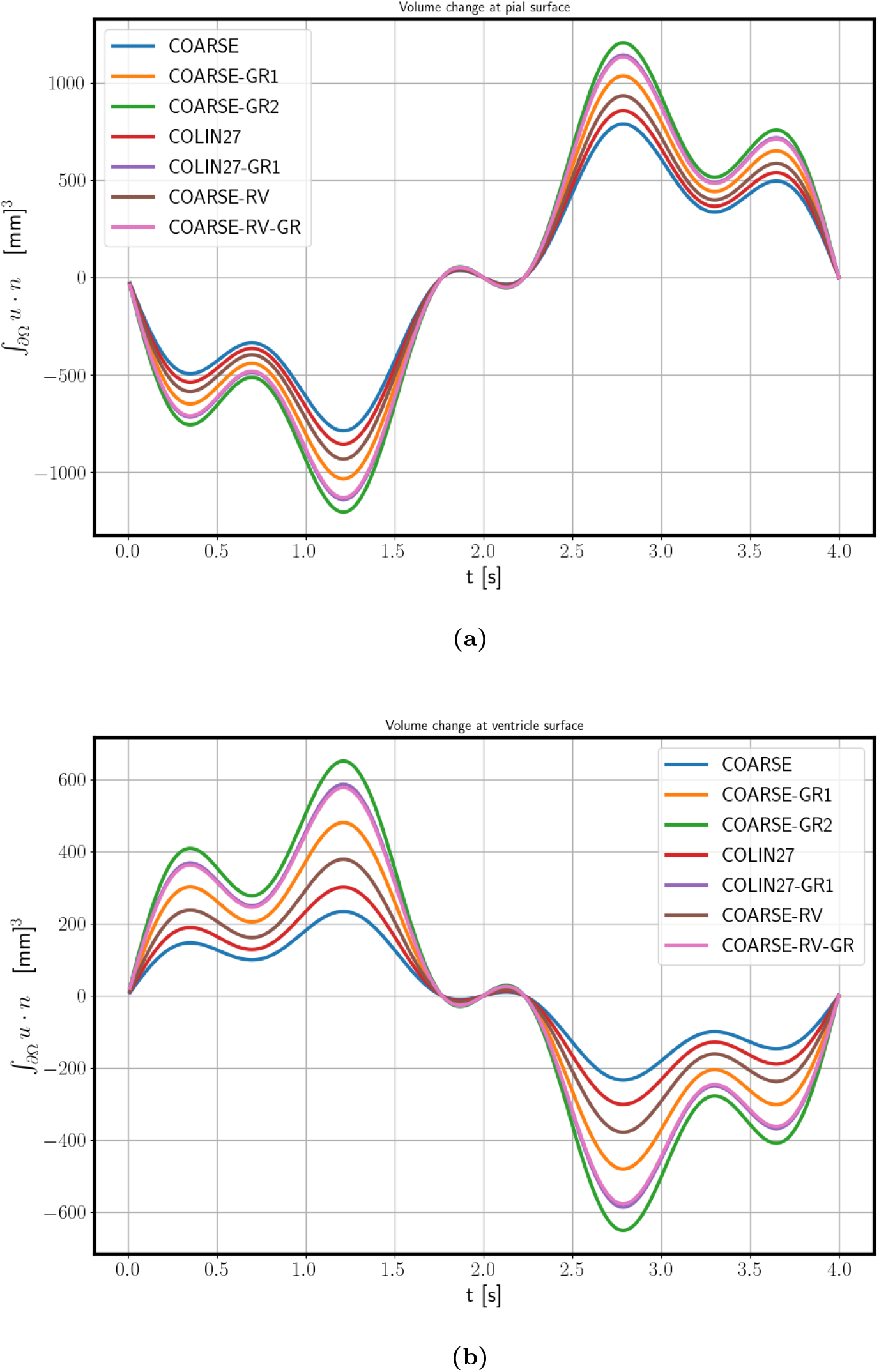
Convergence test for a linear elasticity case over the meshes in Tab. 3: volume change for the pial surface a and for the ventricular surface b. The COARSE-RV-GR-mesh (pink line) was used as the mesh for the numerical simulations.

## Results

### Cardiac pulsatility induces pulsatile brain displacement

The applied pressure difference (Model A) induces a pulsatile displacement of the brain: the parenchyma is initially compressed, and then expands passing through no displacement at *t* = 0.5 s and *t* = 1.0 s. The peak displacement magnitude is 0.15 mm and occurs at *t* = 0.25s and *t* = 0.75s relative to the cardiac cycle (Figure 3). The peak x-, y-, and z-displacements occur at the same times and are 0.10, 0.09 mm and 0.10mm, respectively. The negative and positive displacements are clearly separated, creating an ideal separation line that crosses the ventricle area, where the largest magnitudes are observed (Figure 3).

**Fig 3.**
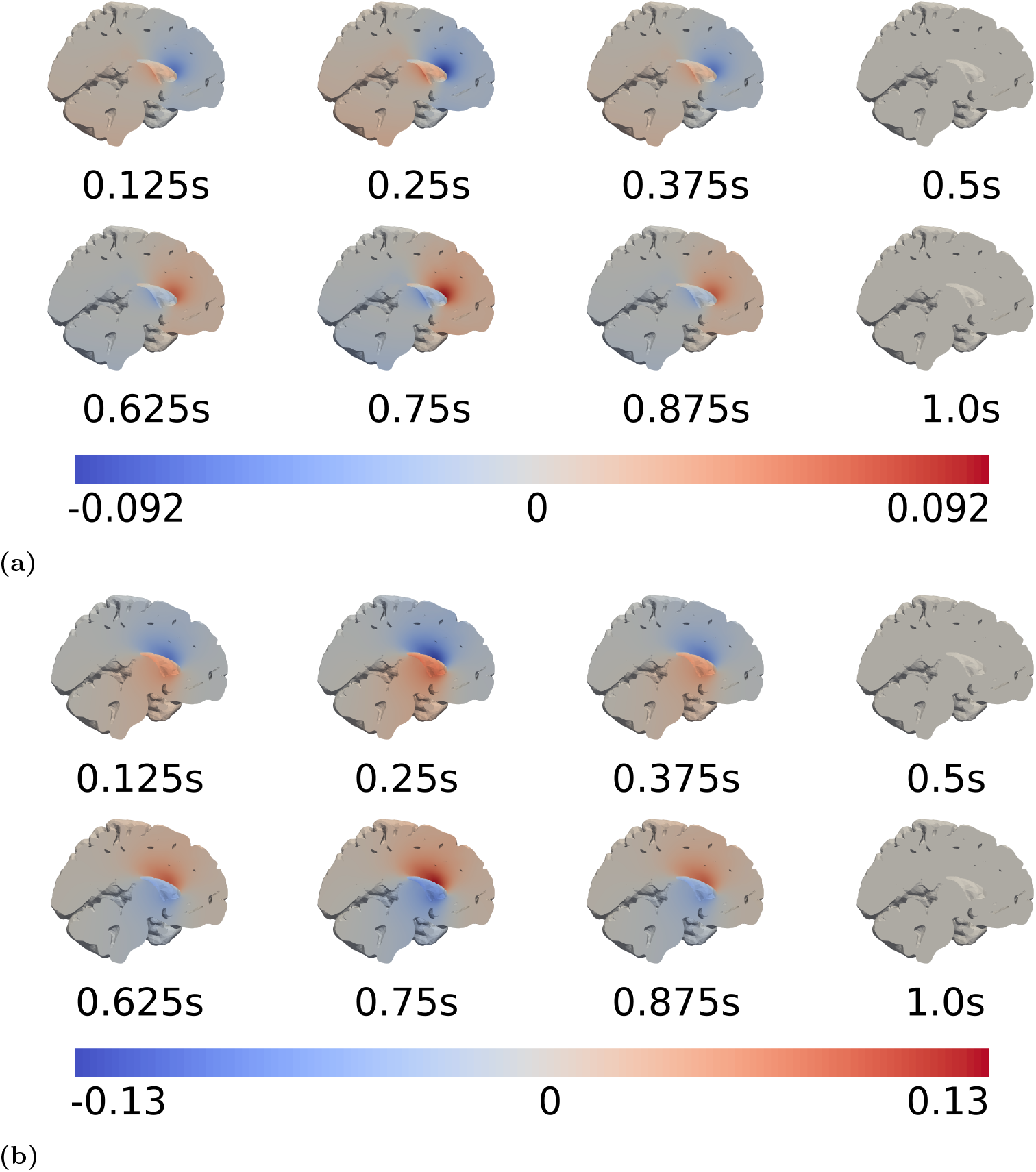
Snapshots at different times relative to the cardiac cycle of the displacement (in mm) induced by a cardiac pressure difference between the pial and ventricular boundary in the y-direction (a), and z-direction (b).

### Cardiac pulsatility dominates respiratory pulsatility in brain displacements

When applying a pressure difference between the pial and ventricular boundaries with both a cardiac and respiratory component (Model B), we observe an analogous behaviour as with only a cardiac contribution but now with a longer period, higher magnitudes, and more local maxima and minima. Again, negative and positive displacements are clearly separated resulting in a ideal separation line that crosses the ventricular area. Again, as the pressure increases on the pial surface (Figure 1) for t < 2 s, the brain is compressed, with a peak compression of 569.87 mm^3^ at 1.25 s and a peak expansion of 569.87 mm^3^ at 2.75s (Figure 4a-b). In particular, the volume change through the pial surface reaches the maximum of 1163.86 mm^3^ at 2, 75s, while the volume change through the ventricular surface reaches the maximum of 594 mm^3^ at 1.25 s. The volume change curves through the parenchyma and through the ventricles (Fig 4a) are shifted by π in phase.

**Fig 4.**
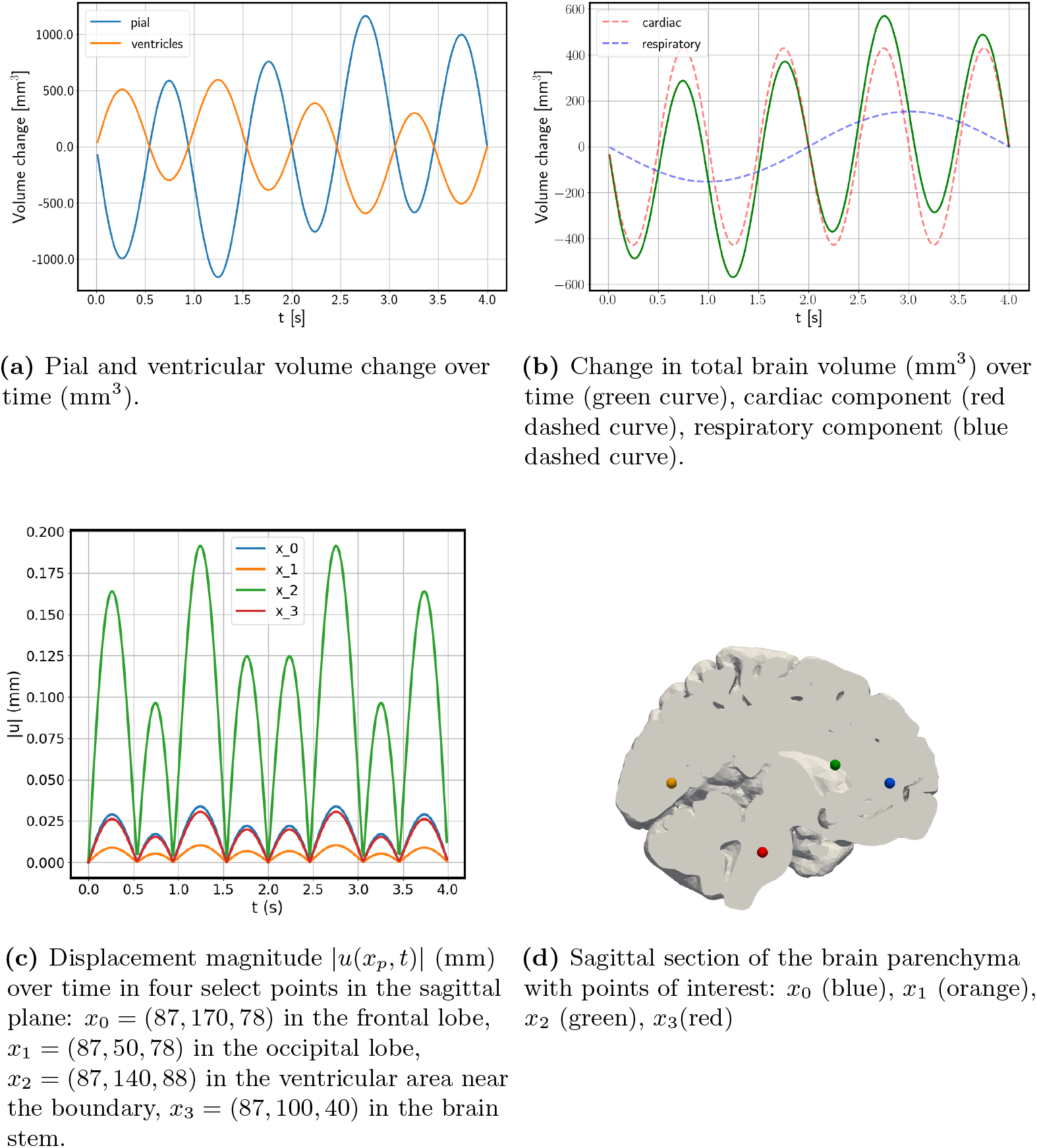
Volume changes and displacement magnitude under cardiac and respiratory pressure pulsations (Model B).

The cardiac component dominates the respiratory component, both in the applied gradient and, as expected, in the induced displacement and volume change (Figure 4b-c). The peak volume change due to the cardiac component of the applied pressure gradient is 429 mm ^3^, while the peak volume change due to the respiratory component of the applied pressure gradient is 153 mm^3^ (Fig 4b). We observe the greatest displacement in the ventricular area (Figure 4c). The peak displacement magnitude max ||*u*|| is 0.196 mm and occurs at *t* = 1.25 s and *t* = 2.75 s above the lateral ventricles near the boundary.

### Brain displacements persist under reduced compressibility

For a more incompressible parenchyma (Model C), we observe the same brain displacement patterns with less than 1.0 % change in displacement magnitude: the peak displacement magnitude for this case is 0.185 mm (compared to 0.196 mm in Model B) (data not shown).

### ISF pressure is nearly uniform with impermeable pial membrane

For the poroelastic case, we also observe an initial compression followed by an expansion of the parenchyma. Comparing with the elastic displacements at *t* = 1.25 s and *t* = 2.75 s, the peak displacement magnitude is 0.22, and occurs in close proximity of the ventricles (Figure 5a). The peak displacement predicted by the poroelastic model is 0.22 mm, which is higher than in the elastic model with the same driving forces. The overall pattern of displacement, including the relative importance of the cardiac versus respiratory component, is similar to displacements observed with the linear elasticity model.

**Fig 5.**
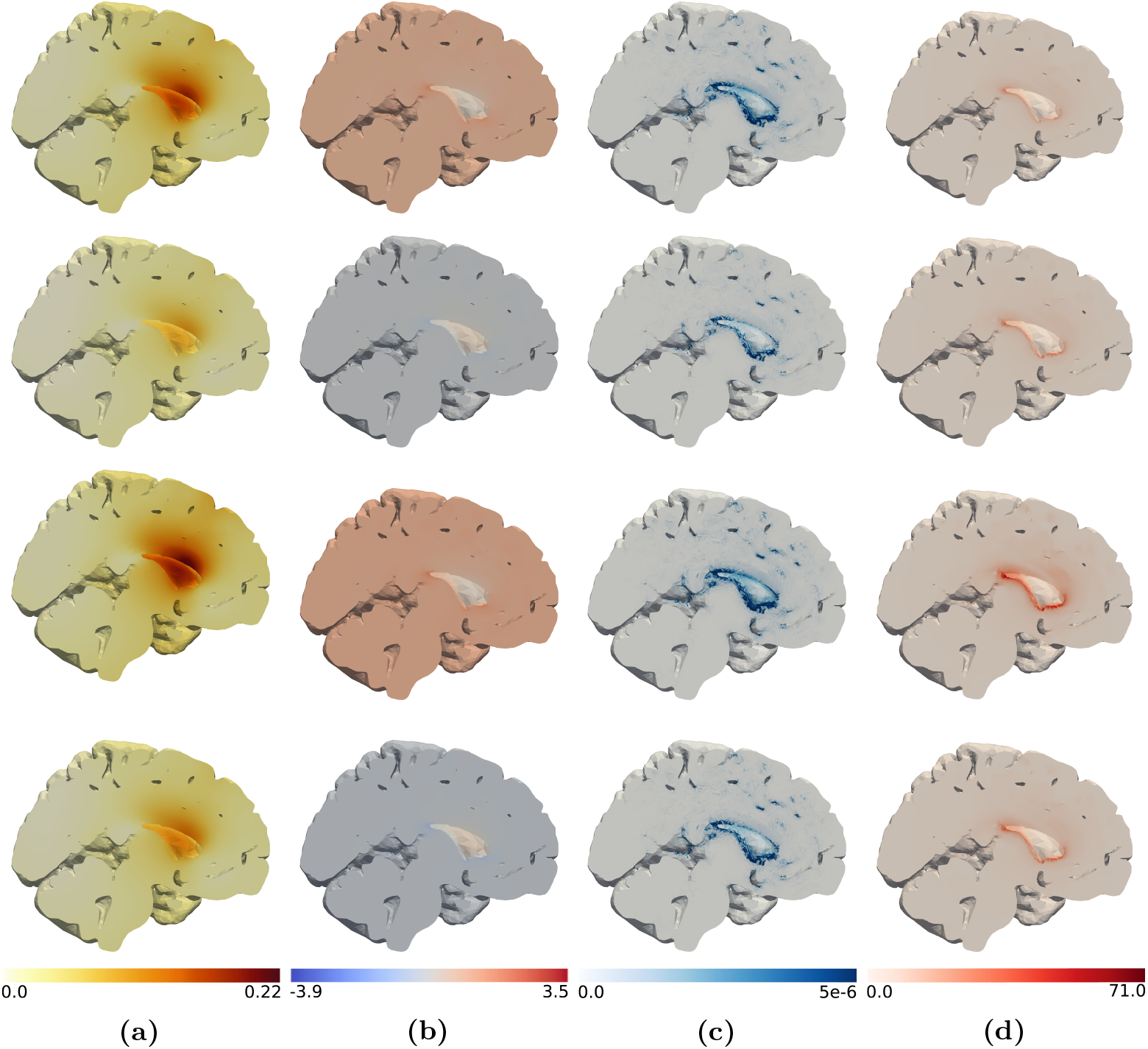
Column (a) displacement magnitude (mm), (b) fluid pressure (Pa), (c) fluid velocity (−*K* ∇*p*) magnitude (mm/s), and (d) von Mises stress (Pa) at different time points: *t* = 0.2625 s, 0.75s, 1.25s, 1.75s (from top to bottom).

During the 4s respiratory cycle, the fluid pressure *p* is nearly uniform in space with minimum and maximum values of −3.9 Pa and 3.5 Pa, respectively (Figure 5b). Again, the extreme values occur in the vicinity of the ventricles. The peak fluid pressure difference is thus lower than the prescribed stress. As the pressure is nearly uniform, the pressure gradient is small almost everywhere in the domain with average pressure gradient magnitude of less than 9.0 · 10^-3^ Pa/mm. In localized regions, high pressure gradients are observed (of up to 16 Pa/mm).

With the pressure gradients reported above, the peak velocity magnitude is 0.26 *μ*m/s and occurs at t = 2.725. The highest velocities occur near the ventricles where the extracellular fluid flows in the same direction as the movement of the ventricular surface induced by the applied pressure difference. In regions further away from the ventricles, fluid velocities are negligible and on the order of a few nm/s.

The solid part of the stress is visualized via the von Mises stress *σ_M_* (Figure 5d). The von Mises stress is nearly zero (< 5Pa) everywhere in the parenchyma, except for in the ventricular area where it reaches its maximum of 134 Pa at 1.25 s. The peak value is only observed in a very limited set of nodes (less than 0.06% of total nodes) in the proximity of the ventricles. In Figure 6 we show the volume change computed at each time step on the pial and on the ventricular surfaces. The volume change for the ventricular and the pial surface are opposite in sign. The maximum volume change magnitudes are reached at 1.2125s and 2.775s and are 764 mm^3^ for the ventricles and 1293 mm^3^ for the pial surface. The total volume change (Fig. 6(b)) reaches its maximum of 529 mm^3^ at 2.775s. From the decomposed signal, we find a cardiac induced stroke volume of 409.86 mm^3^, and a respiratory induced stroke volume of 146.37 mm^3^.

**Fig 6.**
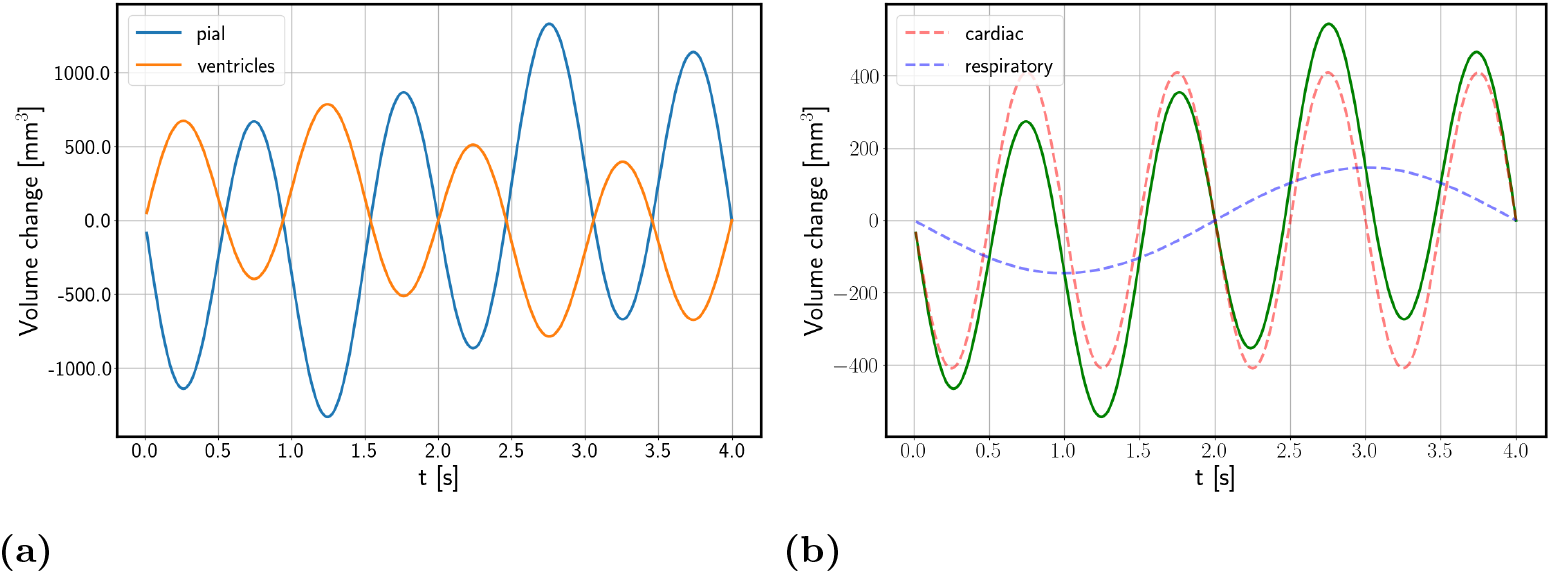
Poroelastic model with impermeable pial membrane driven by cardiac and respiratory pulsatility: (a) volume change on pial (blue) and ventricular (orange) surfaces, (b) total volume change (green curve) and its cardiac component (red dashed curve) and respiratory component (blue dashed curve)

### Pial membrane permeability induces sharp ISF pressure boundary layer

With permeable pial and ventricular membranes (*β* = ∞, model E) the peak displacement magnitude is 0.22 mm at *t* = 2.75s in the ventricular area (data not shown). The characteristics of the displacement field do not change significantly from previous models. The volume change caused by the displacement field is also comparable to what observed with model D (data not shown).

For the pressure *p* (Fig. 7 top) we observe a mostly uniform field but a sharp boundary layer for the pial boundary. The applied pressure gradient and small hydraulic conductivity *K* do not allow for the pulsation to be transmitted inside the parenchymal tissue. We observe the same sharp boundary layer also for the fluid velocity magnitude (Fig. 7). Fig. 8 shows the fluid flux through the ventricular and pial surfaces. We observe that the majority of the flux happens through the pial surface, where it reaches the maximum value of 62.89mm^3^/s, compared to only 0.11 mm^3^/s for the ventricular surface.

**Fig 7.**
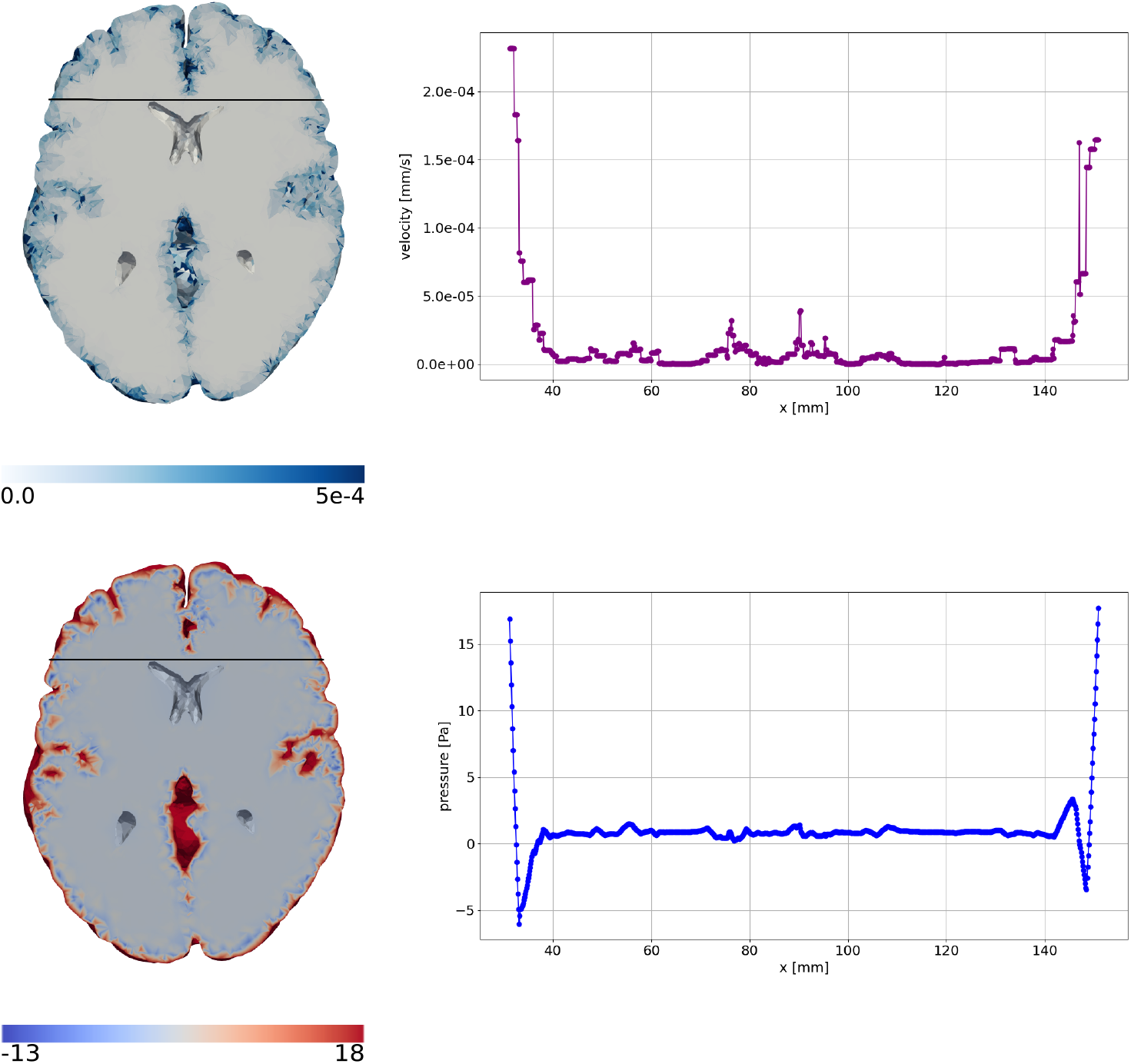
Poroelastic model with fully permeable pial membrane driven by cardiac and respiratory pulsatility: fluid velocity (top left) and fluid pressure (bottom left) in horizontal section. Fluid velocity magnitude (mm/s) over the black line on the horizontal section(top right), pressure (Pa) over the black line on the horizontal section(bottom right).

**Fig 8.**
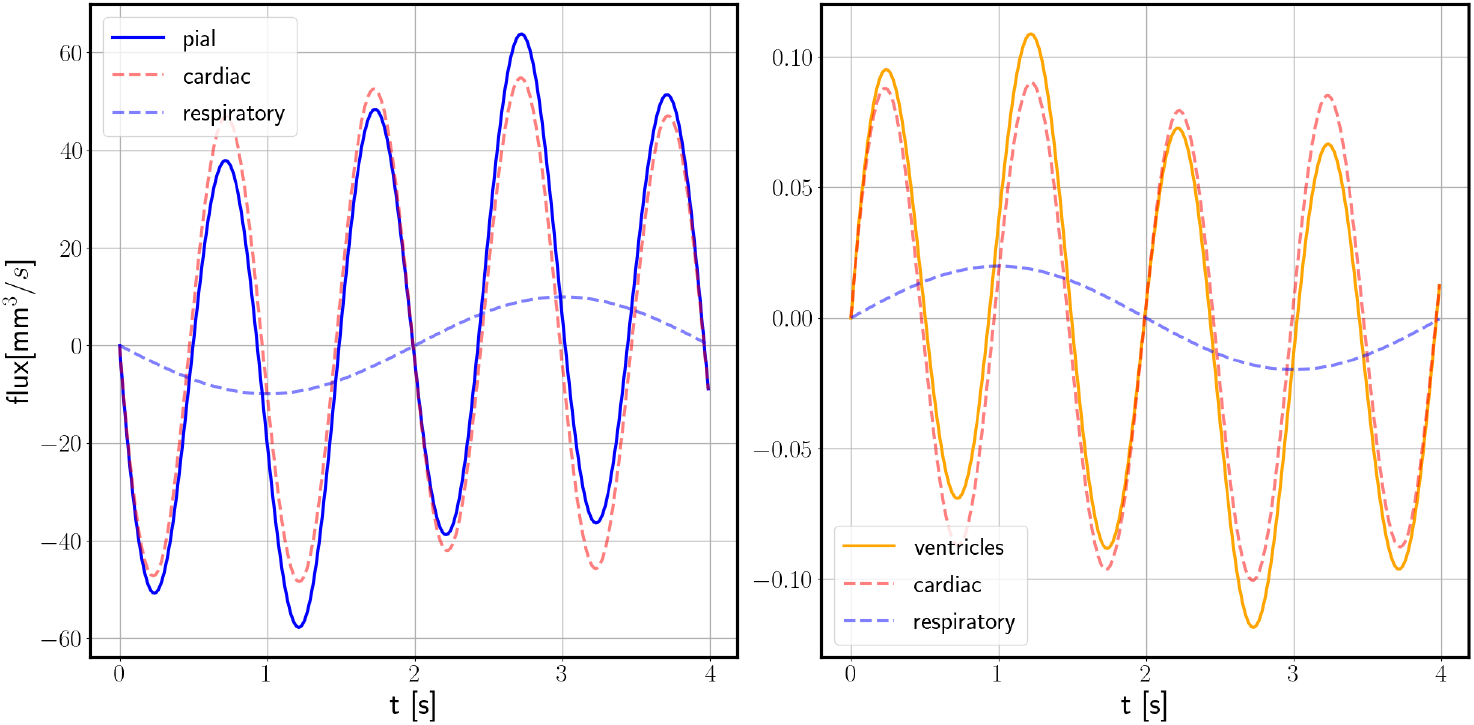
Poroelastic model with fully permeable pial membrane driven by cardiac and respiratory pulsatility: fluid flux on pial (blue solid curve) and ventricular (orange solid curve) surfaces. Red and blue dashed curves show the cardiac and respiratory components.

From the decomposed signal (Figure 8), we find an average cardiac peak volumetric flux of 54.79 mm^3^/s for the pial and 0.09 mm^3^/s for the ventricular surface. The corresponding peak respiratory fluxes are lower, 9.95 mm^3^/s for the pial and 0.028 mm^3^/s for the ventricular surface. Integrating the individual curves (Figure 8) gives a cardiac-induced volume change of 8.19 mm^3^ for the pial and 0.015 mm^3^ for the ventricular surface. The corresponding respiratory-induced volume changes are 6.335 mm^3^ for the pial and 0.01 mm^3^ for the ventricular surface.

## Discussion

### Summary of results

The importance of cardiac versus respiratory pulsations on brain displacements are not yet fully understood. In this study, we have shown that in vivo measurements of pressure differences within the cranium [9] induce pulsatile brain displacements with both cardiac and respiratory components. Even the more basic linear elastic model provides useful insights regarding the displacement of parenchyma. In fact, the difference between the displacement fields obtained with the linear elastic model (B) and the poroelastic models (D and E) are negligible. Furthermore, the cardiac pulsation alone is responsible for the largest part of the displacements occurring in the brain parenchyma. For the poroelastic models, the impermeable boundary condition (model D) results in a mostly uniform pressure field, and therefore almost zero, and mostly concentrated in the ventricular area, fluid velocity. The fully permeable boundary condition (model E), results in a sharp boundary layer both for the pressure and on the velocity field. The pulsation is not transmitted through the fluid in the parenchymal tissue but it remains on the domain boundary. This behaviour suggests that, with the given, physiological, material parameters, a systemic pressure gradient alone is not sufficient to drive fluid movement through the brain.

### Comparison with literature

The maximal displacement magnitude (with models D and E) is 0.22 mm, in excellent agreement with values for peak displacement reported by Pahlavian et al. and Sloots et al. [24, 25]. Assuming that these displacements occur over a segment of approximately 6 cm, the maximal volumetric strain is 3.3 × 10^-3^, which is exactly what was measured by Sloots et al. [25]. Pahlavian et al. [24] pointed to medial and inferior brain regions as regions with large motion, while our model predicts peak displacements in regions in close proximity to the ventricles (Figure 3). In addition, the experimental values reported in [24, 25] only took into account motion induced by cardiac pulsations, while the respiratory influence was overlooked. In our model, the applied pressure pulsation induced by the cardiac cycle is 2-3 times larger than the respiratory pulsation, and a similar relationship is obtained for the displacements induced by the two cycles. The peak displacement induced solely by the cardiac pulsation reached 0.15 mm, comparable to values in [24]. It is worth noting that this linear relationship between pressure and movement will not necessarily hold for CSF flow in the SAS [10]. A higher value for the von Mises stresses near the ventricles in our simulations suggests that this region is most prone to shape distortion caused by the cardiac and respiratory cycles. As reported in preliminary work by Sincomb and colleagues [26,27], due to the viscoelastic nature of the brain, even a small transmantle pressure gradient can over time contribute to the enlargement of the ventricles. The viscoelastic nature of the brain may explain why some authors have assumed the brain to be relatively compressible when modeling long-term behaviour (e.g. hydrocephalus) [16,28,29].

As brain and CSF movement is coupled, changes in brain volume, as computed by our models, will result in CSF flow in the SAS. It is reasonable to assume that flow and displacements on the ventricular surface is directly related to aqueductal flow, while flow and displacements on the pial surface may be related to flow in the foramen magnum. The total volume change peaks (models B, D, and E) and the respective stroke volumes associated with the cardiac component are comparable. We found a cardiac induced stroke volume of 429*μ*L (model B), and 529*μ*L (model D) compared to approximately 500*μ*L at C2-C3 in [30] (Table 11.1 for healthy subjects). On the other hand, the volume change through the ventricles estimated from model B is 593*μ*L, and is approximately one order of magnitude larger than the ventricular stroke volumes reported for healthy subjects 48*μ*L [30] Table 11.1). Furthermore, the aqueductal stroke volume computed with all models (linear elasticity and poroelasticity) is closer to the reported values for idiopathic normal pressure hydrocephalus patients [30] Table 11.1). An observed delay in the reversal of flow in the cerebral aqueduct compared to the foramen magnum [31] was not predicted by volumetric changes in our model.

ISF flow within the human brain has not been measured experimentally, but several estimates have been made. From experimental data of tracer distribution and clearance in rats, Cserr and colleagues estimated a bulk flow velocity of around 0.1–0.25 *μ*m/s [32, 33]. A directional bulk flow of this magnitude in addition to diffusion may explain tracer movement in humans [34]. On the brain surface of mice, pulsatile CSF flow of magnitudes of around 20 *μ*m/s have been observed on top of a static flow of similar magnitude [13,35]. Flow of ISF in our model occurred mainly close to the pial surface and a peak velocity of 0.5*μ*m/s (model E) were observed. Fluid flow within the parenchyma was dominated by cardiac pulsations, contributing a factor 3 more than respiration to fluid flow velocities. The fluid exchange between ISF and CSF (stroke volume induced by fluid flow) for the pial was computed to be 8.19*μ*L over the cardiac and 6.34*μ*L over the respiratory cycle. The fluid exchange over the ventricles is negligible: 0.015*μ*L for the cardiac component, and 0.013*μ*L for the respiratory component. The amount of CSF/ISF-exchange was thus equally dominated by cardiac and respiratory pulsations as the respiratory pulsation spans over a longer time. Several other studies have pointed to respiration to be the main driver of displacement of fluid in the SAS [10, 36, 37]. However, relationships between fluid pressure and flow will differ between CSF and ISF, and it is not given that the pulsations within the parenchyma found in our model translates directly to the CSF in the SAS.

### Limitations

We considered homogeneous properties for the parenchyma tissue without distinguishing between gray and white matter and we modelled the parenchyma tissue as isotropic. Nevertheless, the estimated values for white and gray matter Young modulus are similar [19], and the average value can be used as a good approximation without affecting the results significantly. Moreover, Budday and colleagues [38] demonstrated that the brain tissue can be considered as an isotropic material from a mechanical point of view despite being anisotropic.

In this work, we considered a limited set of parameter values based on the literature currently available. The main goal of this paper was not to perform a parametric study but rather study the effect of a pulsatile pressure gradient on the brain parenchyma. Certainly, a parametric study, taking into account further parameter combinations, could be performed in later work.

The brain tissue is permeated by several fluid networks: ISF, capillary blood, venous blood and arterial blood [2]. In this work, we considered a one–network poroelastic model and we did not consider the exchange between the ISF and the other compartments. The interaction between ISF and other fluid compartments could be modelled with a multiple–network poroelastic model [16,29] and it could be investigated in future work.

The CSF fluid dynamics in the ventricles and in subarachnoid space was not included in our models. In particular, the resistance to flow through the aqueduct, SAS or spinal canal is not explicitly modeled The resistance is probably much higher in aqueduct, which may explain why our models overestimate aqueductal flow (volume change through the ventricular surface), but not flow to the spinal flow (volume change through the pial surface).

Finally, we note that the pressure data was obtained from iNPH patients [10], which may serve as another source of error, particularly for aqueductal stroke volume [30]. However, to our knowledge no such intracranial in-vivo pressure measurements exists from healthy volunteers.

### Conclusion

We have presented elastic and poroelastic models of the brain with pulsatile motion driven by pressure pulsations originating from the cardiac and respiratory cycle. The displacement fields and total volume change match well with values found in the literature, while the pressure applied on the boundary did not properly propagate through brain tissue, suggesting that pressure pulsations from blood vessels act not only on the surface, but also within brain tissue. Further investigation of pressure pulse propagation within the brain parenchyma is needed to fully understand the mechanisms leading and connected to brain parenchyma pulsation and brain clearance.

